# GDA-Pred: Generative AI-Driven Data Augmentation for Improved Prediction of IL-6 and IL-13 Inducing Peptides

**DOI:** 10.64898/2025.12.07.692883

**Authors:** Hiroyuki Kurata, Hiroto Tsuruta, Soyogu Shigetomi, Md. HARUN-OR-ROSHID, Kazuhiro Maeda

## Abstract

The identification of interleukin-6 (IL-6) and interleukin-13 (IL-13) inducing peptides is crucial for accelerating drug discovery targeting cancer, immune disorders, and infectious diseases. However, experimental screening methods remain time-consuming and costly. To address these limitations, various machine learning and deep learning models have been developed, yet their performance is still constrained by the limited availability of experimentally validated data. In this study, we propose a generative AI-driven data augmentation (GDA) framework and predictor, GDA-Pred, to improve the prediction performance of state-of-the-art (SOTA) classifiers for identifying IL-6 and IL-13 inducing peptides. GDA expands the training dataset by generating novel peptide sequences using three types of generative AI models: generative adversarial networks (GANs), diffusion models (DMs), and variational autoencoders (VAEs). The GDA framework is defined by four key parameters: the type of generative model, the sequence identity cutoff, the probability threshold (PT) for selecting generated peptides, and the augmentation ratio (AR) between generated and real peptides. Since optimizing these parameters is challenging with small datasets, we adopt a case study-oriented proof-of-concept approach using a moderately sized dataset of anti-inflammatory peptides (AIPs) to derive interpretable optimal settings. The performance of the optimized GDA was evaluated using stratified 5-fold cross-validation with cluster-based partitioning and a hold-out test on benchmark datasets. The optimized GDA was then applied to SOTA classifiers, collectively termed GDA-Pred, to identify IL-6 and IL-13 inducing peptides, both of which are limited by small dataset sizes. GDA-Pred substantially improved the prediction performance for both cytokine-inducing peptide tasks, demonstrating the feasibility of GDA-Pred as a robust and generalizable framework.

## 1. Introduction

Functional peptides, defined as short amino acid sequences with specific biological activities, have emerged as a promising class of therapeutic agents in modern drug discovery [1–3]. Unlike small-molecule drugs, which often exhibit limited specificity and off-target effects, peptides can closely mimic endogenous ligands and protein–protein interaction motifs, enabling highly selective binding to cellular receptors, enzymes, and other molecular targets. This high specificity not only enhances therapeutic efficacy but also reduces adverse side effects, leading to improved safety profiles in clinical applications. Their functional versatility allows them to act as anticancer, antimicrobial, antivirus, anti-immune, cytokine-inducing, hormonal, and neuromodulatory peptides, thereby expanding the range of treatable diseases.

Within this landscape, cytokine-inducing peptides, which elicit the secretion or modulation of specific cytokines from immune cells, represent a promising avenue for advanced immunomodulatory drug discovery or immune-modifying therapeutics by selectively engaging innate or adaptive immune pathways. These offer a level of control not easily achieved with small molecules or broadly neutralizing antibodies. Interleukin-6 (IL-6)-inducing and interleukin-13 (IL-13)-inducing peptides exemplify complementary therapeutic peptides. They share common principles in cytokine regulation and act through receptor-mediated signaling pathways, influence downstream transcription factors, and modulate innate and adaptive immunity [4–7]. Since IL-6 is a pleiotropic mediator central to both innate and adaptive immunity, IL-6-inducing peptides predominantly elicit pro-inflammatory responses and are associated with acute-phase reactions, T helper 17 (Th17) cell differentiation, and autoimmune and inflammatory conditions such as rheumatoid arthritis, IgA nephropathy, and the cytokine storm associated with severe COVID-19 [4, 5, 8, 9]. In contrast, since IL-13 is a canonical T-Helper 2 (Th-2) cytokine, IL-13–inducing peptides drive Th-2-mediated immune responses, contributing to allergic inflammation, parasitic infections, mucus production, and differentiation of naive human B cells toward IgG4- and IgE-producing cells [6, 7]. Peptide-based vaccines and peptide kinoids targeting IL-13 demonstrated preclinical efficacy in attenuating airway inflammation and allergic phenotypes [10].

Despite these potentials, experimental identification of these peptides is time-intensive and resource demanding. Computational methods for their prediction and characterization are critically required, because it enables rapid and cost-effective screening of vast sequence spaces that would be impractical to evaluate experimentally [2, 11, 12]. To date machine learning (ML)-and deep learning (DL)-based methods have been intensively proposed for identifying IL-6-inducing peptides [13–18] and for IL-13-inducing peptides [19–21], while publicly available datasets for these cytokine classes remain extremely limited [15, 19, 20]. They implemented a broad range of ML and DL architectures in combination with diverse peptide encodings, including composition, position-order and physicochemical-based property representations, and language model embeddings such as Evolutionary Scale Modeling-2 (ESM-2) [22]. Kurata et al. [21] and Harun-Or-Roshid et [17] al. proposed an advanced ensemble learning framework in which a logistic regression model and genetic algorithms integrate the probability scores from over 100 individual ML and DL models, each trained on a single feature representation, respectively. Cao et al developed a graph neural network model that incorporate 3D structure information [23] to identify IL-6-inducing peptides. However, the prediction performance of these models remains suboptimal, primarily due to the extremely limited availability of public datasets for these cytokine classes [15, 19, 20]. There are few data augmentation studies that challenge the scarcity of experimentally validated cytokine-inducing peptides[24].

To date many data augmentation methods had been developed including Synthetic Minority Oversampling Technique (SMOTE) [25] and Kernel Density Estimation (KDE) [26]. In peptide sequence analysis, data augmentation methods such as ACP-DA[27] and ACP-ADA [28] were employed to address the limited dataset of anticancer peptides (ACPs). ACP-DA encoded peptide sequences using a combination of binary profile features and AAindex-derived physicochemical properties, and generated new samples in the feature space to enrich the training dataset [27]. ACP-ADA employed an AdaBoost-based framework that integrates binary profile features, amino acid indices, and amino acid composition [28].

These days generative AIs including Generative adversarial networks (GANs) [29, 30], diffusion models (DMs)[31, 32], and variational autoencoders (VAEs) [33] are expected as alternatives to conventional data augmentation methods. Such AI-based data augmentations, named generative AI-driven data augmentation (GDA), would learn the underlying distribution of known functional peptides and generate diverse and informative representation of the sequence space while preserving key physicochemical and functional properties. Augmenting the training set with these high-quality generated, fake peptides would increase the prediction performance of classifier models. However, generative AI–based data augmentation is still in its early stages, and only a limited number of studies have explored its potential to improve predictive performance. For example, Deep convolutional GANs effectively expand training datasets by generating ACPs, particularly enriching underrepresented classes [29]. Such GDA strategies remain insufficiently established and there are few guidelines for their implementation. It is required to present some intelligible guidelines for GDA, including criteria for choosing model architectures of AIs, selecting high-quality generated peptides, and determining an optimal augmentation ratio of the generated peptides to the training dataset.

To address the limitations arising from the small dataset sizes of IL-6- and IL-13-inducing peptides, we have developed a robust GDA framework that expands the peptide sequence space by incorporating AI-generated peptides into the training dataset. The GDA framework was optimized using a moderately sized dataset of anti-inflammatory peptides (AIPs) and subsequently integrated into state-of-the-art (SOTA) predictive models, termed GDA-Preds, for the identification of IL-6- and IL-13-inducing peptides. This development followed a case study-oriented proof-of-concept approach, in which AIPs provided an interpretable example, as the limited size of cytokine-inducing peptide datasets poses significant challenges for GDA parameter optimization.

## 2. Methodology

### 2.1. Development flow

As illustrated in **Figure 1**, the development of GDA-Pred comprises four main steps: (1) Generate fake peptide sequences from the real, positive peptides of the training dataset by generative AIs. (2) Select the generated peptides via a classifier built with the training dataset. (3) Add the generated peptides to the training dataset. (4) Construct classification predictors with the expanded training dataset and optimize key GDA parameters, (5) Evaluate the prediction performance of the optimized GDA-based classifiers (GDA-Pred). The four key parameters are: (i) the type of generative AI (GAN, DM, or VAE); (ii) the sequence identity cutoff value in the Needleman–Wunsch algorithm (NW) [34] for redundancy removal; (iii) the probability threshold (PT) for selecting generated peptides; and (iv) the augmentation ratio (AR) of generated peptides added to the training dataset.

**Figure 1.**
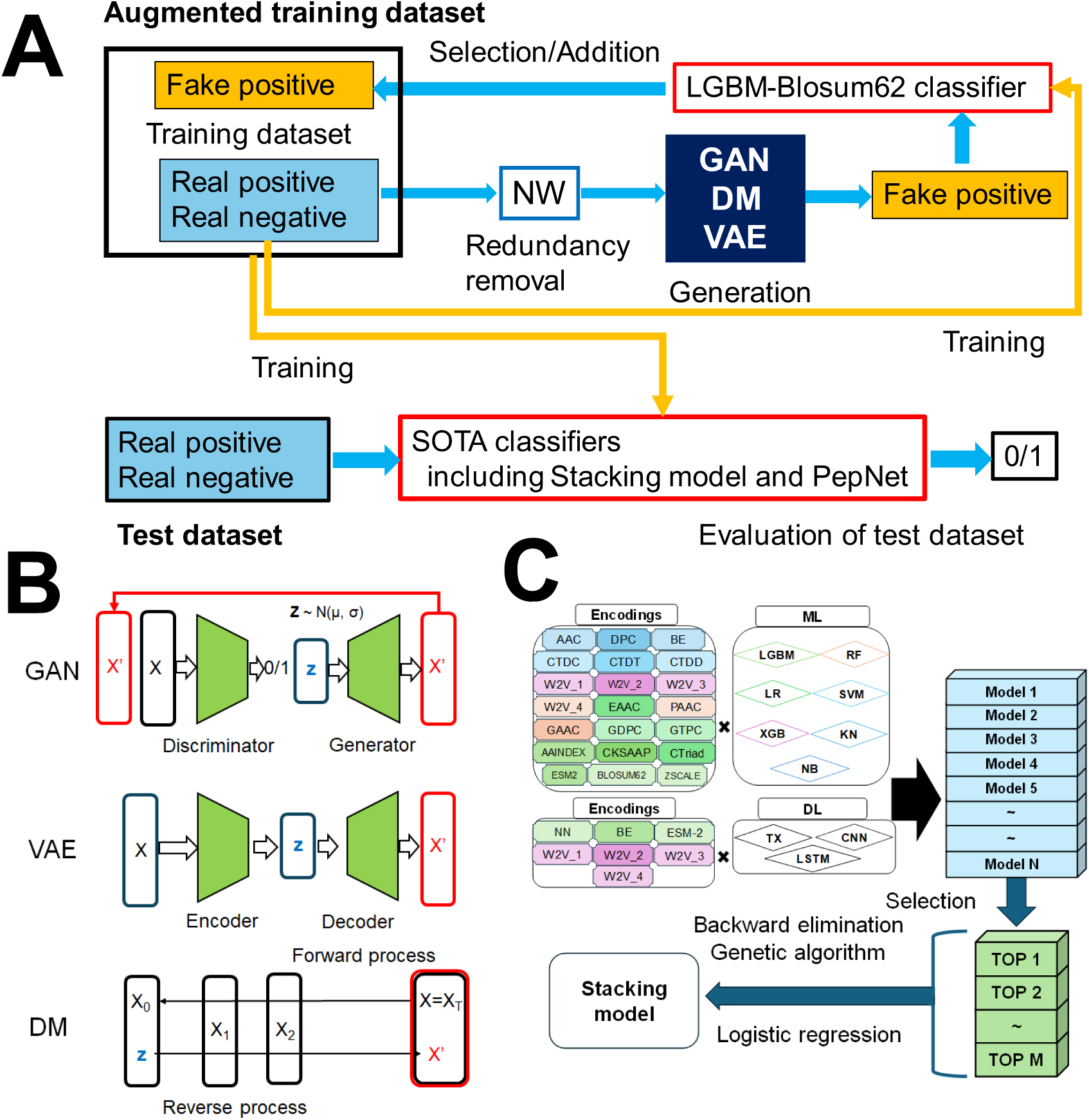
Development Workflow of GDA-Pred. (A) GDA framework consisting of (1) Generate fake peptide sequences from the real, positive peptides of the training dataset by generative AIs. (2) Select the generated peptides by a classifier constructed with the training dataset. (3) Add the generated peptides to the training dataset. (4) Construct classification predictors with the expanded training dataset, (5) Test the predictors with an independent test dataset. (B) Architectures of three generative AIs: GAN, DM, and VAE. (C) Our stacking classifiers for prediction of functional peptides.

### 2.2. Benchmark datasets

A curated dataset of experimentally validated bioactive peptide sequences was obtained from publicly available peptide databases [15, 19, 20, 24, 35], as shown in **Table 1**. Sequences were filtered to include only peptides with experimentally confirmed activity to exclude non-standard amino acids, and limited to a specified length range (e.g., 5–50 residues).

**Table 1.** Benchmark datasets.

| Peptide | Training dataset |  | Test dataset |  |
| --- | --- | --- | --- | --- |
|  | Positive | Negative | Positive | Negative |
| IL6-inducing peptide | 292 | 2393 | 73 | 598 |
| IL13-inducing peptide | 250 | 2326 | 63 | 582 |
| AIP | 1258 | 1887 | 420 | 629 |

### 2.3. Cluster-based 5-fold partitioned datasets

Redundant sequences (i.e., highly similar or identical peptides) can bias generative models to overfit some common patterns and cause data leakage. To eliminate redundant peptide sequences from the benchmark dataset, we employed the Needleman–Wunsch (NW) algorithm [34]-based redundancy removal method (NW-RR) using a sequence identity cutoff of 0.4, 0.5, 0.6, 0.7, 0.8, and 0.9 (**Figure S1**). The sequences were clustered based on their similarity such that only one representative sequence from each cluster was retained.

The non-redundant datasets are divided into 5-fold balanced and diverse clusters, as follows (**Figure S1**). We employed a clustering approach based on pairwise alignment similarity. First, all-against-all global pairwise alignment scores between peptide sequences were computed using the NW algorithm implemented in Biopython’s pairwise2 module [36], with standard scoring parameters (match = +1, mismatch = −1, gap open = −10, gap extend = −0.5). The resulting similarity matrix was transformed into a dissimilarity matrix by subtracting each score from the maximum observed score. To reduce the high-dimensional similarity space, we applied classical multidimensional scaling (MDS) to project the sequences into a two-dimensional Euclidean space. Subsequently, k-means clustering (k = 5) was performed on the MDS coordinates to partition sequences into five groups. To ensure balanced dataset sizes, we randomly down-sampled each cluster to match the size of the smallest cluster. To validate the generalizability of GDA, we employed the cluster-based 5-fold partitioned datasets, where samples were grouped into five balanced and diverse clusters for 5-fold cross-validation (CV).

### 2.4. Generation of peptide sequences by generative AIs

Three generative AIs (GAN, DM, VAE) are used to generate fake, positive peptide sequences from the real, positive sequences of the non-redundant training dataset processed with NW-RR. Peptide sequences are encoded by giving 6 physicochemical property indexes to each peptide residue. Output vectors are decoded into approximate residue vectors using cosine similarity. Details of model architectures and tuning parameters are shown in **Figure 1** and **Table 2**.

**Table 2.**
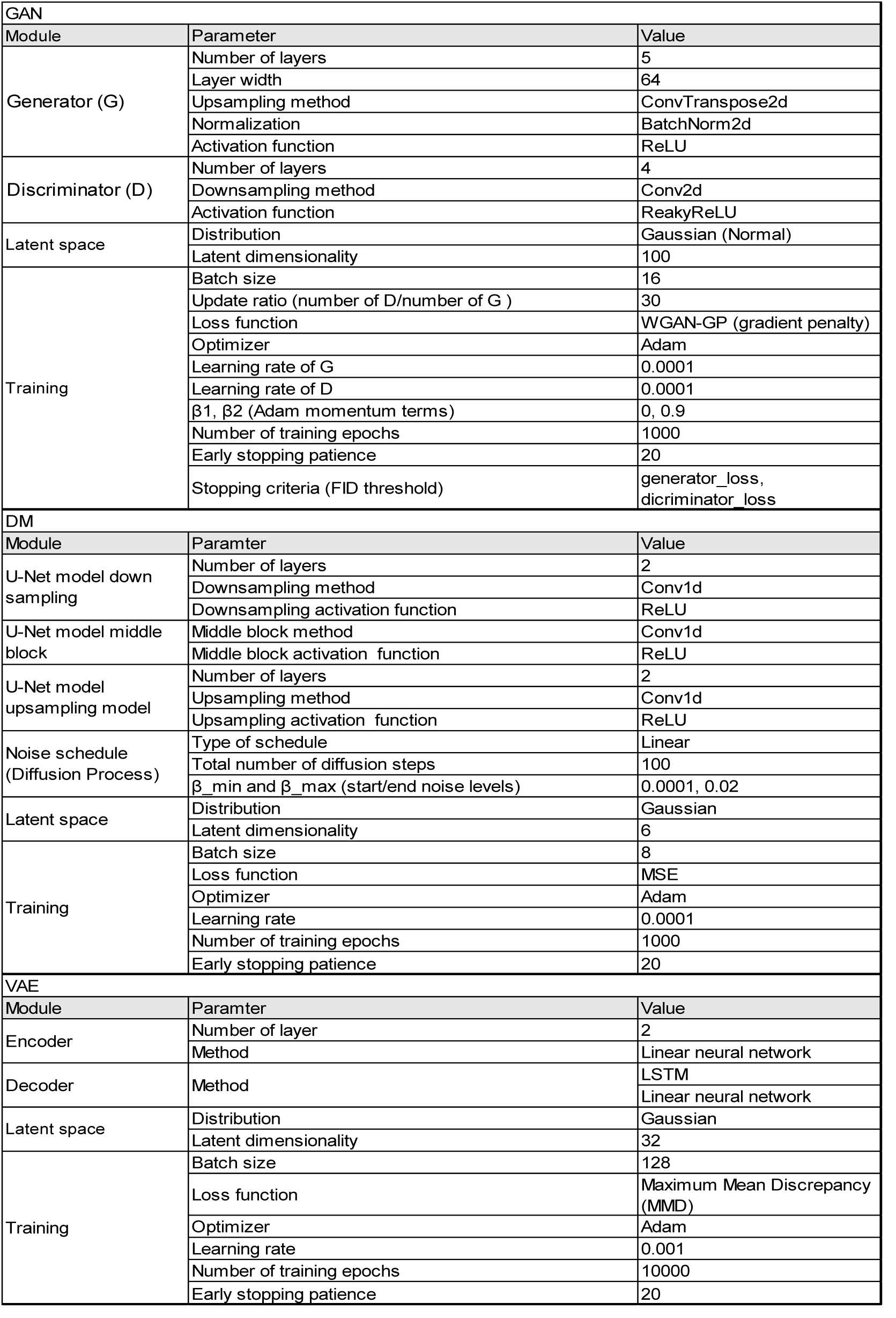
Details of model architectures and tuning parameters for the three generative AIs (GANs, DMs, and VAEs)

#### GAN Architecture

Based on the previous work [30], we built a GAN consisting of Two-Dimensional Convolutional Neural Network (2DCNN)-based generator and discriminator. These were trained in a minimax game framework with real positive sequences of functional peptides. The generator takes random noise vectors as input and generates fake peptide sequences. The discriminator receives real peptide sequences (from the dataset) and generated sequences and outputs a probability score indicating whether the sequence is real or fake.

#### DM architecture

Based on [31, 32], we built a DM model to learn the generative distribution of functional peptides. The forward process gradually added Gaussian noise to the peptide embeddings over a predefined number of time steps. The reverse process was parameterized using a U-Net comprising CNN layers to predict and remove the noise at each step.

#### VAE architecture

To learn a compact latent representation of the peptide sequences, we implemented a sequence-to-sequence VAE model [33]. The encoder consisted of stacked neural network that encoded the input peptide sequence into a fixed-length latent vector. The decoder, also based on Long Short Term Memory (LSTM) layers, reconstructed the peptide sequence from the latent representation. Once the autoencoder is trained, novel peptide sequences are generated by sampling latent vectors from the learned latent space.

### 2.5. Conventional data augmentation method

To generate positive sequences, we used two conventional data augmentation methods: Synthetic Minority Oversampling Technique (SMOTE) [25] and Kernel Density Estimation (KDE) [26] as control methods. SMOTE identifies its *k* nearest neighbors in the feature space for each positive sequence. After a random neighbor is selected, a new sequence is generated along the line segment connecting the original sequence and its neighbor (*k = 5*). KDE is a non-parametric method for estimating the underlying probability distribution of a dataset. It estimates the data distribution by summing kernel functions, typically Gaussian kernels, centered at each data point. After fitting the KDE model with a bandwidth of 1.0 to the original data, new peptide sequences are generated by sampling from the estimated distribution.

### 2.6. Augmentation of training dataset

To assess the biological relevance of the generated peptides, we employed the Light Gradient Boosting Machine (LGBM) classifier with the BLOSUM62 encoding [37, 38], trained with the real positive and negative sequences alone, to give a probability score to each generated peptide (**Figure 1**). We selected the generated peptides with a probability of more than PT as the fake peptides, and then added them to the training dataset with an AR, defined as the ratio of selected peptides to the number of positive training peptides.

### 2.7. Baseline classifiers

We constructed 18 baseline models, the LGBM models combined with 18 different features, and evaluated the prediction performance of the models with the training and test datasets [17, 21, 35]. LGBM stands as a gradient boosting platform harnessing tree-based learning algorithms, presenting heightened efficiency and swiftness. The distinctive features of LGBM encompass adept handling of expansive datasets, superior effectiveness, and rapid execution. What distinguishes it is the employment of a histogram-based algorithm, discretizing continuous feature values into distinct bins, a departure from conventional tree-based methods.

The amino acid sequences were encoded using 18 different encoding techniques including Amino Acid Composition (AAC), Di-Peptide Composition (DPC) [39], Composition of k-spaced Amino Acid Pairs (CKSAAP)[40], Grouped Amino Acid Composition (GAAC) [41], Grouped Dipeptide Composition (GDPC), Grouped Tripeptide Composition (GTPC), Composition/Transition/Distribution (CTD) Composition (CTDC), CTD Transition (CTDT), CTD Distribution (CTDD) [42, 43], Binary Encoding (BE), Enhanced Amino Acid Composition (EAAC)[44], Amino Acid indices (AAindex) [45], BLOSUM62 [46], and, Word2Vec (W2V) with different Kmers of 1, 2, 3 and 4 [47]. All these sequence encoding techniques can be readily computed using the open-source software packages including iLearn [44]. These encoding techniques capture a variety of properties including compositional, position-order, evolutional and physicochemical property, and linguistic patterns/distributions. Each of these encoding methodologies brings a unique lens to the sequence and contributes to a comprehensive and detailed analysis.

### 2.8. SOTA classifiers

As SOTA models, we employed PredIL6 [17], PredIL13 [21], and PepNet [48] (**Figure 1**). In PredIL6 and PredIL13, seven ML algorithms (LGBM, eXtreme Gradient Boosting, random forest, support vector machine, logistic regression, naïve bayse, and k-nearest neighbor) and three DL architectures (convolutional neural network, attention, and LSTM) were integrated with a wide range of sequence encoding techniques, resulting in construction of hundreds of baseline models. The predicted probabilities from these baseline models were combined using a logistic regression-and genetic algorithm-based stacking approach [17, 21]. The optimized stacking model was evaluated via 5-fold CV on the benchmark training dataset to ensure that the predicted probabilities were well calibrated with the true class labels. PepNet was developed as an interpretable neural network for predicting both AIPs and anti-microbial peptides (AMPs) by applying a pre-trained protein language model [48]. PepNet captures the peptide sequence information of residue arrangements and physicochemical properties using a residual dilated convolution block and then obtains function-related diverse information by introducing a residual transformer block that characterizes the residue representations generated by the pre-trained protein language model. The GDA-Preds, the SOTA classifiers trained with the GDA-expanded benchmark training dataset, were evaluated on the benchmark test dataset.

### 2.9. Quality of generated peptides

To examine the quality of generated (fake) peptides, we calculated the dipeptide frequency distribution and 8 physicochemical properties including Aliphatic Index (Thermal Stability) [49], Aromaticity (Aromatic AA Content) [50], Boman Index (Molecular Compactness) [51], Net charge, Charge Density (Charge per Length) [50], Hydrophobic Ratio (Hydrophobicity) [52], Instability Index (Atmospheric Stability), Isoelectric Point (Acid-Base Property) [53]. To visualize high-dimensional data of peptide sequence embeddings on a 2D map, Uniform Manifold Approximation and Projection (UMAP) is used as dimensionality reduction techniques [54].

### 2.10. Measures

Prediction performance was assessed using 6 established statistical measures, each offering different insights into the prediction performance [55]. It includes accuracy (ACC), sensitivity (SEN), specificity (SPE), precision (PRE), Matthews correlation coefficient (MCC), and area under the receiver operating characteristic curve (AUC). Comprehensive descriptions and mathematical formulations of these measures are available in previous studies. A threshold value that determines whether the probability scores are classified into positive and negative samples is adjusted so as to maximize MCC.

To make clear the effectiveness of GDA, the performance increase ratio (PIR) is defined by:

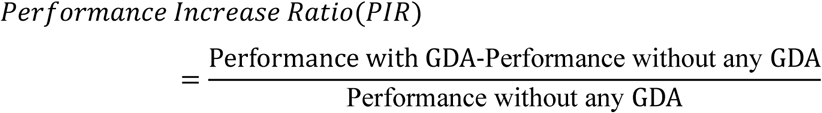

where performance without any GDA is named control performance. A positive value of PIR indicates that GDA increases classification performance; a negative value that GDA decreases it.

All programs were implemented in Python using Biopython [36], scikit-learn libraries[56] and PyTorch [57].

## 3. Results

### 3.1. Peptide generation

The GAN model was trained using non-redundant, real positive sequences from the AIP training dataset, which were preprocessed with NW-RR using a similarity cutoff of 0.7. The GAN subsequently generated positive peptide sequences from Gaussian noise. The quality of the generated peptides was evaluated, as shown in **Figure 2**. The generated peptides exhibited slightly higher median values of aliphatic index, net charge, charge density, hydrophobic ratio and isoelectric point compared with the real peptides, whereas the remaining physicochemical properties were largely similar between the two groups. Although the compositions of cysteine (C) and tryptophan (W) differed, those of the other amino acids were comparable. The UMAP visualization of the embedding vectors derived from physicochemical property indices revealed substantial overlap between the distributions of the generated and real peptides. These findings suggest that the generated sequences relatively resemble the real positive sequences in both physicochemical characteristics and sequence patterns. The generated peptides via DMs and VAEs also exhibited similar properties to the real ones, as shown in **Figures S2** and **S3**.

**Figure 2.**
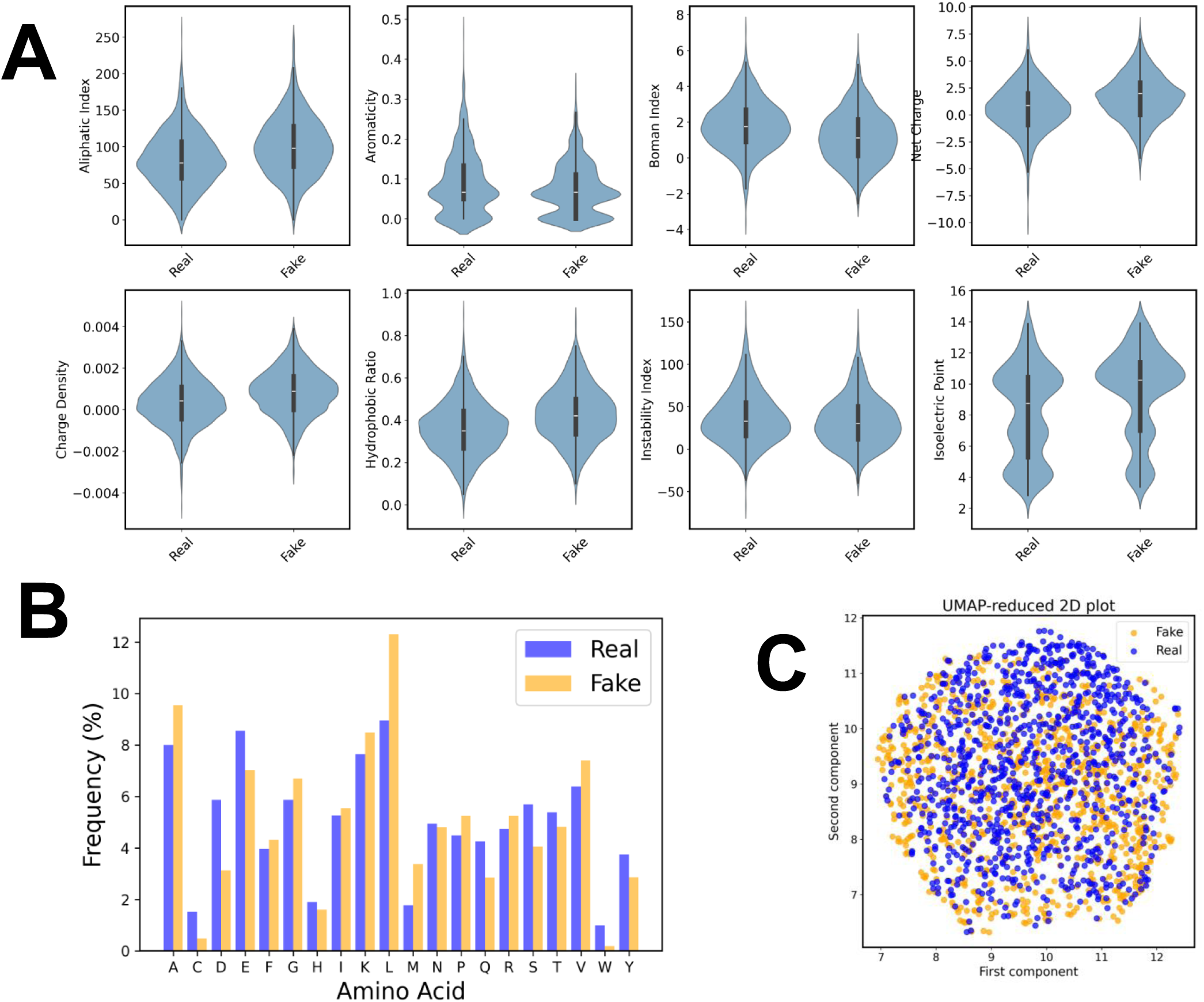
Evaluation of the quality of AIP fake peptide sequences that GANs generate from the real, positive peptides of the training dataset. The fake peptides correspond to the generated peptides. The fake AIPs are generated from the non-redundant training dataset built with NW-RR with a sequence identity cutoff of 0.7. (A) Physicochemical properties. (B) Frequency distribution of amino acid residues. (C) UMAP distributions of the encoding vectors of real and fake peptide sequences.

### 3.2. Stratified 5-fold CV with cluster-based partitioning

The generated peptides were evaluated by an LGBM model with the BLOSUM62 encoding trained with the AIP training dataset, to assign probability scores to them. We selected the generated peptides with probability values greater than 0.5 (PT) as augmentation data. To investigate the generalizability of the GDA, we added the selected AIP peptides to the training dataset with an AR of 0.25, trained the 18 baseline models with the expanded training dataset, and evaluated the models via stratified 5-fold CV with cluster-based partitioning (**Figure S4**). To highlight the effectiveness of GDA, the corresponding PIR values of AUC, MCC, and ACC are illustrated in **Figure 3**. In the training dataset, applying GDA enhanced the AUC, MCC, and ACC of all baseline models across every fold. Although the test dataset exhibited lower performance due to strict sequence similarity-based cluster partitioning, GDA improved AUC, MCC, and ACC in all but 4 of the 90 cases (18 encodings × 5 folds). Application of GDA distinctly increased the generalizability of AIP prediction

**Figure 3.**
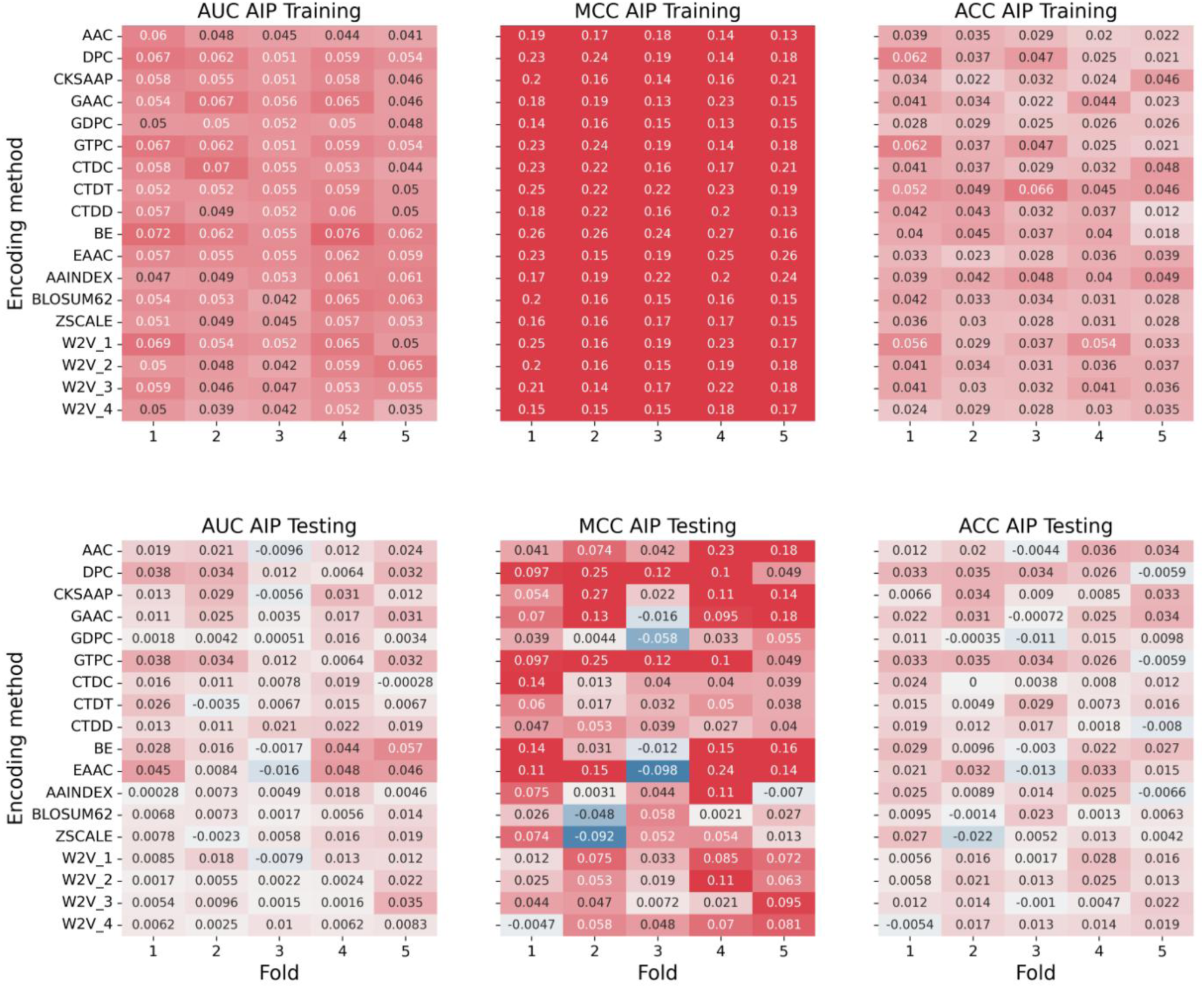
PIRs of AUC, MCC, and ACC for 18 baseline classifiers trained by the augmented training dataset with GAN-generated AIPs via the stratified 5-fold CV with cluster partitioning. Fake AIP sequences were generated using GANs from a non-redundant training dataset built by NW-RR with a sequence identity cutoff of 0.7. The generated peptides were filtered using an LGBM classifier with BLOSUM62 encoding at a PT of 0.5, and the selected sequences were added to the training dataset with an AR of 0.25. Red and blue colors indicate an increase and a decrease in PIRs, respectively.

### 3.3. Optimality of GDA

In **Figure S4** and **Figure 3**, we presented the GDA method with four optimized key parameters: the type of GAN, a cutoff of 0.7, a PT of 0.5, and an AR of 0.25. Here, we demonstrate the optimality of these four parameters. First, we investigated which generative AI was most suitable for GDA using a cutoff of 0.7, an AR of 0.25, and a PT of 0.5. **Figure S5** shows that the prediction performances of 18 baseline classifiers trained with the augmented dataset by GAN, DM and VAE via stratified 5-fold CV with cluster-based partitioning. In the training dataset, GAN and DM increased the PIR values of MCC for all the baseline classifiers across all 90 cases (18 classifiers x 5 folds), while VAE presented negative PIR values of MCC in several cases. In the test dataset, GAN increased the PIR value of MCC more than DM in many cases, while VAE showed less PIRs of MCC than GAN and DM. Among the three generative models, GAN achieved the best performance, exhibiting slightly higher scores than DM and substantially higher than VAE.

Second, we analyzed how a cutoff of NW-RR influenced the performances of 18 baseline models trained with the GAN-expanded datasets. Using different cutoffs of 0.5, 0.6, 0.7, 0.8 and 0.9, we evaluated the AUC, MCC and ACC values the 18 baseline classifiers with an AR of 0.25 and PT of 0.5 via stratified 5-fold CV with cluster-based partitioning, as shown in **Figure S6**. The AUC, MCC and ACC remained a robust property across a wide range of a cutoff from 0.5 to 0.9 on both the training and test datasets with respect to baseline models. Based on these results, the cutoff value was set to 0.7 in this study.

Third, we investigated how a PT value altered the performances of 18 baseline models trained with the GAN-expanded datasets. Using different PT values of 0.0, 0.5 and 0.7, we evaluated the PIR values of MCC for the baseline classifiers with a cutoff of 0.7 and an AR of 0.25 via stratified 5-fold CV with cluster-based partitioning, as shown in **Figure S7**. In the training dataset, the PIR values of most of baseline classifiers across all the folds increased with increasing PT, reaching saturation at a PT of 0.5. The GDA improved the MCC values for all baseline models. In contrast, in the test dataset, the number of the baseline classifiers exhibiting positive PIR values increased with PT across all folds, peaked at PT of 0.5, and then declined at a PT of 0.7. At a PT of 0.7, overfitting was observed. Therefore, a PT of 0.5 was identified as the optimal threshold for model generalization. These results indicate that a PT of 0.5 provided the optimal balance, resulting in an appropriately diverse sequence distribution. In contrast, peptides generated with a PT of 0.0 exhibited excessive diversity, whereas those generated with a PT of 0.7 showed restricted diversity.

Fourth, we examined the effect of an AR on prediction performance of 18 baseline models trained with the expanded datasets. Using AR values of 0.1, 0.25, and 0.4, we evaluated the PIR values of MCC for the baseline classifiers with a cutoff of 0.7 and a PT of 0.5 via stratified 5-fold CV with cluster-based partitioning, as shown in **Figure S8**. In the training dataset, the PIR values of the baseline classifiers across all folds increased progressively with the AR, attaining the highest values at an AR of 0.4. In contrast, in the test dataset, the number of the baseline classifiers exhibiting positive PIR values increased with AR across all folds, reached a peak at an AR of 0.25, and subsequently declined. At an AR of 0.4, overfitting was observed. Accordingly, the optimal AR was determined to be 0.25.

Finally, we applied the GDA method to the benchmark datasets of AIPs (**Table 1**). After removing redundancy from the benchmark training dataset with a NW-RR cutoff of 0.7, GAN was used to generate fake AIP sequences. The benchmark training dataset was augmented with an AR of 0.2 by using the selected, generated peptides with a PT of 0.5. We trained our stacking classifier (PredIL6) and the PepNet with the expanded training dataset, and evaluated the models on the test dataset, as shown in **Table 3**. Applying GDA to both PredIL6 and PepNet substantially increased most of statistical measures compared to the control method, whereas conventional data augmentation methods (SMOTE and KDE) reduced SEN, ACC, MCC, and AUC. The integration of GDA improved their performances with respective increases of 0.4% and 0.5% in AUC, 1.2% and 1.3% in MCC, and 0.3% and 1.3% in ACC for the stacking classifier and PepNet. PepNet exhibited higher scores of SEN, PRE, ACC, MCC, and AUC than the stacking model, probably because PepNet was specifically developed for the identification of AIPs [48].y

**Table 3.** GDA-enhanced prediction performances of SOTAs for identifying AIPs on the benchmark AIP test dataset. Two SOTA models were employed to predict AIPs with and without applying GDA. GDA is performed using GANs with a cutoff of 0.7, a PT of 0.5, and an AR of 0.25.

| Peptide | Classifier | DA | SEN | SPE | PRE | ACC | MCC | AUC |
| --- | --- | --- | --- | --- | --- | --- | --- | --- |
| AIP | PredIL6 | GDA | **0.627** | **0.790** | **0.670** | **0.725** | **0.424** | **0.766** |
|  |  | SMOTE | 0.394 | 0.837 | 0.625 | 0.660 | 0.263 | 0.696 |
|  |  | KDE | 0.334 | 0.893 | 0.686 | 0.669 | 0.281 | 0.724 |
|  |  | w/o | 0.624 | 0.789 | 0.665 | 0.723 | 0.419 | 0.763 |
|  | PepNet | GDA | 0.708 | **0.831** | **0.738** | **0.782** | **0.546** | **0.841** |
|  |  | SMOTE | 0.678 | 0.818 | 0.715 | 0.762 | 0.501 | 0.817 |
|  |  | KDE | 0.679 | 0.800 | 0.697 | 0.752 | 0.483 | 0.801 |
|  |  | w/o | **0.760** | 0.780 | 0.702 | 0.772 | 0.539 | 0.837 |

### 3.4. Effect of dataset size on GDA

To examine how a training dataset size affects the effectiveness of GDA, the full benchmark dataset of AIPs was divided into training and test sets, while setting the ratio of the positive and negative data to be 1 and the five training datasets of different sizes were prepared. We then applied the optimized GDA method to the AIP datasets with sizes of 160, 320, 640, 1280 and 2560 and evaluated the PIRs of AUC, MCC, and ACC on both the training and test sets, as shown in **Figure 4**. At a small dataset size of 160, several baseline classifiers exhibited negative PIR values in both the training and test datasets, indicating that GDA did not improve prediction performance. This suggests that the classifiers were unable to fully capture sequence features due to the limited number of peptides. At dataset sizes of 320 and 640, GDA enhanced the prediction performance of all baseline models on the training dataset and increased the number of the baseline models exhibiting positive PIR values in the test dataset. It indicates that GDA increases the generalizability. At larger dataset sizes of 1280 and 2560, GDA improved the prediction performance of all the baseline models in the training dataset; however, the PIR values of many classifiers decreased in the test dataset, indicating reduced generalizability.

**Figure 4.**
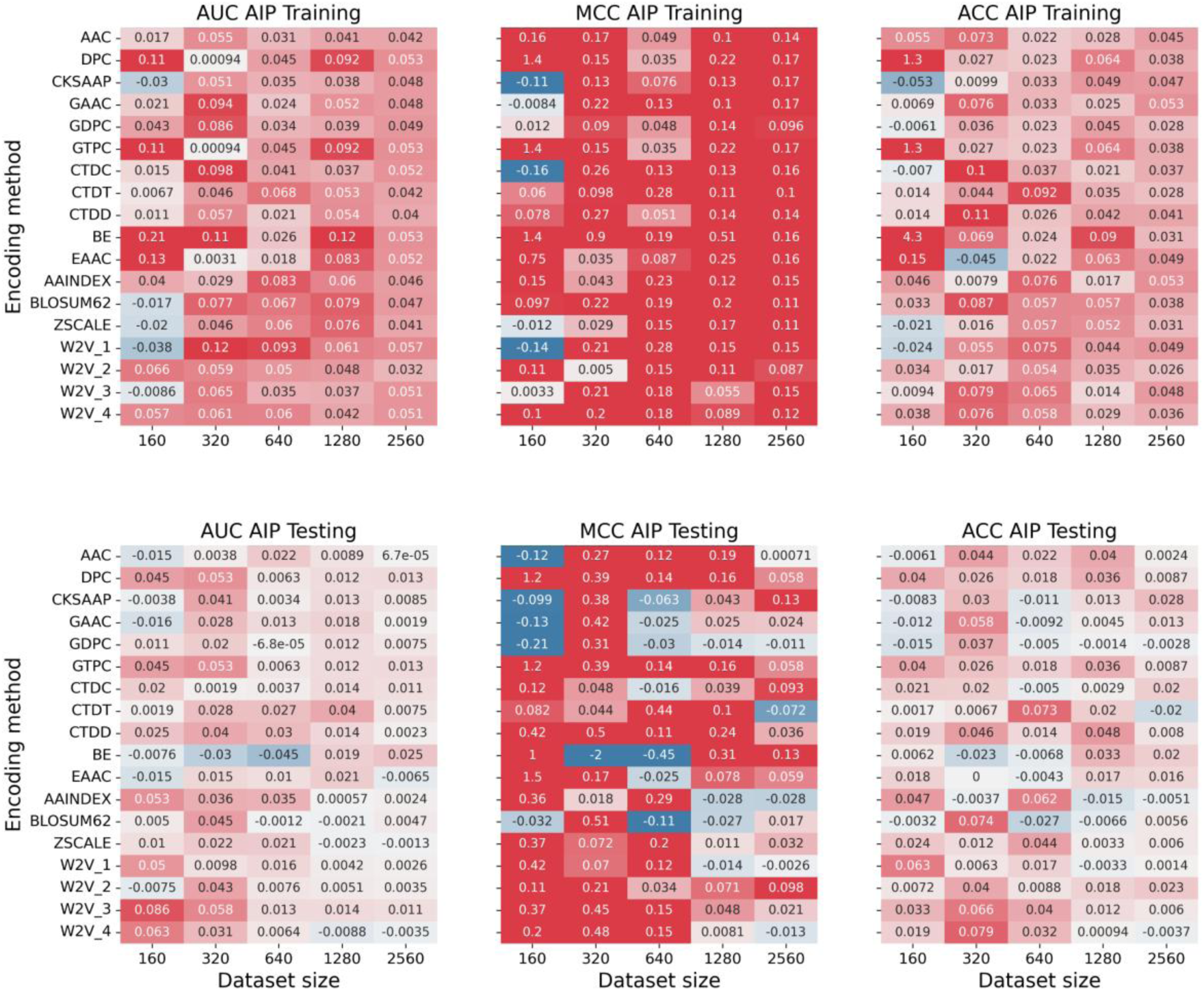
Effect of a training dataset size on PIRs of AUC, MCC, and ACC for 18 baseline classifiers trained by the augmented training dataset with GAN-generated AIPs via 5-fold CV. The optimal parameter settings of GDA-GAN were a cutoff of 0.7, an AR of 0.25, and a PT of 0.5.

We found that GDA was most beneficial when the dataset is moderate or limited, where new generated data can fill gaps in the distribution. When the training dataset is small, GDA cannot capture the true data distribution. It easily overfits to a few samples or cover only a narrow region of the sequence space. When the dataset is very large, the generative model learns a rich and comprehensive representation of the sequences, thus, additional generated data may not provide meaningful new information, which makes augmentation ineffective.

### 3.5. GDA application to IL-6- and IL-13-inducing peptides

Following the optimization of the GDA method parameters by using AIPs, we applied the optimized GDA method to identification of IL-6- and IL-13-inducing peptides. The qualities of the GAN-generated IL-6- and IL-13-inducing peptides derived from the benchmark training dataset were evaluated, as shown in **Figures 5, 6**. For IL-6–inducing peptides, the overall distributions of physicochemical properties between the generated and real peptides were broadly similar. However, noticeable differences were observed in the distribution shapes of aromaticity, hydrophobic ratio, and isoelectric point, as well as in the median values of aromaticity, Boman index, net charge, charge density, hydrophobic ratio, instability index, and isoelectric point. The amino acid composition analysis revealed that the frequencies of glutamic acid (E), phenylalanine (F), methionine (M), proline (P), arginine (R), tryptophan (W) differed between the generated and real peptides, whereas those of the remaining residues were comparable. For IL-13–inducing peptides, the distributions of most physicochemical properties between the generated and real peptides were similar except for aromaticity, and their median values were also comparable except for the instability index. The amino acid frequencies were largely consistent between the generated and real peptides, except for proline (P). Furthermore, UMAP visualization based on physicochemical property embeddings showed comparable distributions between the generated and real peptides for both IL-6 and IL-13 datasets. In summary, the GAN-generated IL-6–inducing peptides exhibited moderately similar sequence and property patterns to the real peptides; the generated IL-13–inducing peptides demonstrated a high degree of similarity to their real counterparts.

**Figure 5.**
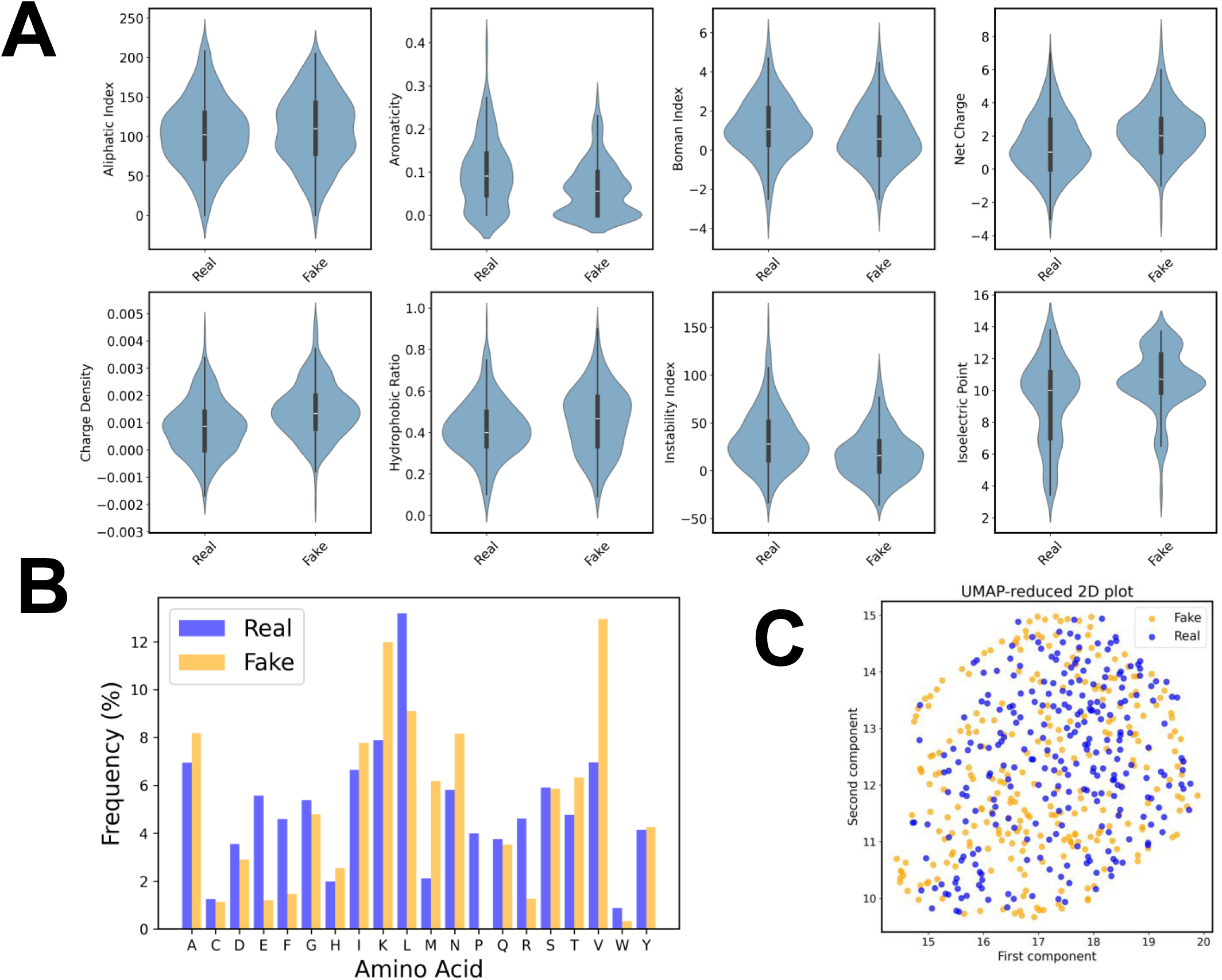
Evaluation of the quality of generated IL6-inducing peptide sequences that GANs generate from the real, positive peptides of the training dataset. The IL-6-inducing peptides are generated from the non-redundant training dataset built with NW-RR with a sequence identity cutoff of 0.7. (A) Physicochemical properties. (B) Frequency distribution of amino acid residues. (C) UMAP distributions of the encoding vectors of real and fake peptide sequences.

**Figure 6.**
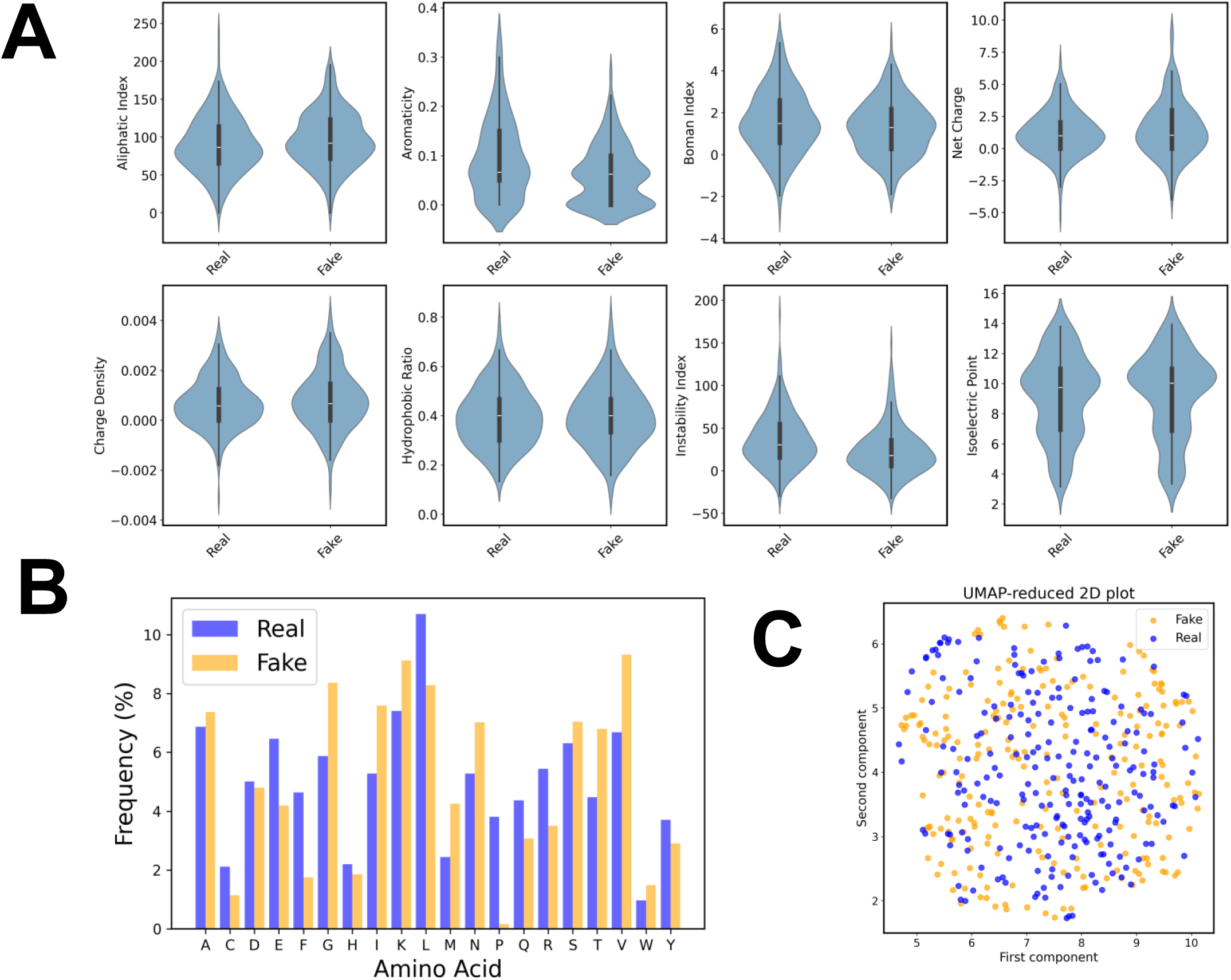
Evaluation of the quality of generated IL-13-inducing peptide sequences that GANs generate from the real, positive peptides of the training dataset. The IL-13-inducing peptides are generated from the non-redundant training dataset built with NW-RR with a sequence identity cutoff of 0.7. (A) Physicochemical properties. (B) Frequency distribution of amino acid residues. (C) UMAP distributions of the encoding vectors of real and fake peptide sequences.

To demonstrate the effectiveness of GDA in identifying IL-6- and IL-13-inducing peptides, we trained the 18 baseline models with the GAN-augmented training dataset with a cutoff of 0.7, a PT of 0.5 and an AR of 0.25 and evaluated them via stratified 5-fold CV with cluster-based partitioning. **Figures 7, 8** illustrate the PIRs of AUC, MCC, and ACC on the training and validation datasets. Applying GDA enhanced the AUC, MCC, and ACC of all the baseline models across every fold on the training dataset for both the cytokine inducing peptides. The test dataset presented lower performance than the training one due to rigorous testing with cluster partitioning. In IL-6-inducing peptides, GDA improved AUC, MCC, and ACC in all but 27 of the 90 cases (18 encodings × 5 folds). In IL-13-inducing peptides, GDA improved AUC, MCC, and ACC in all but 16 of the 90 cases (18 encodings × 5 folds). The GDA substantially increased the generalizability of both the cytokine-inducing peptide prediction in many cases.

**Figure 7.**
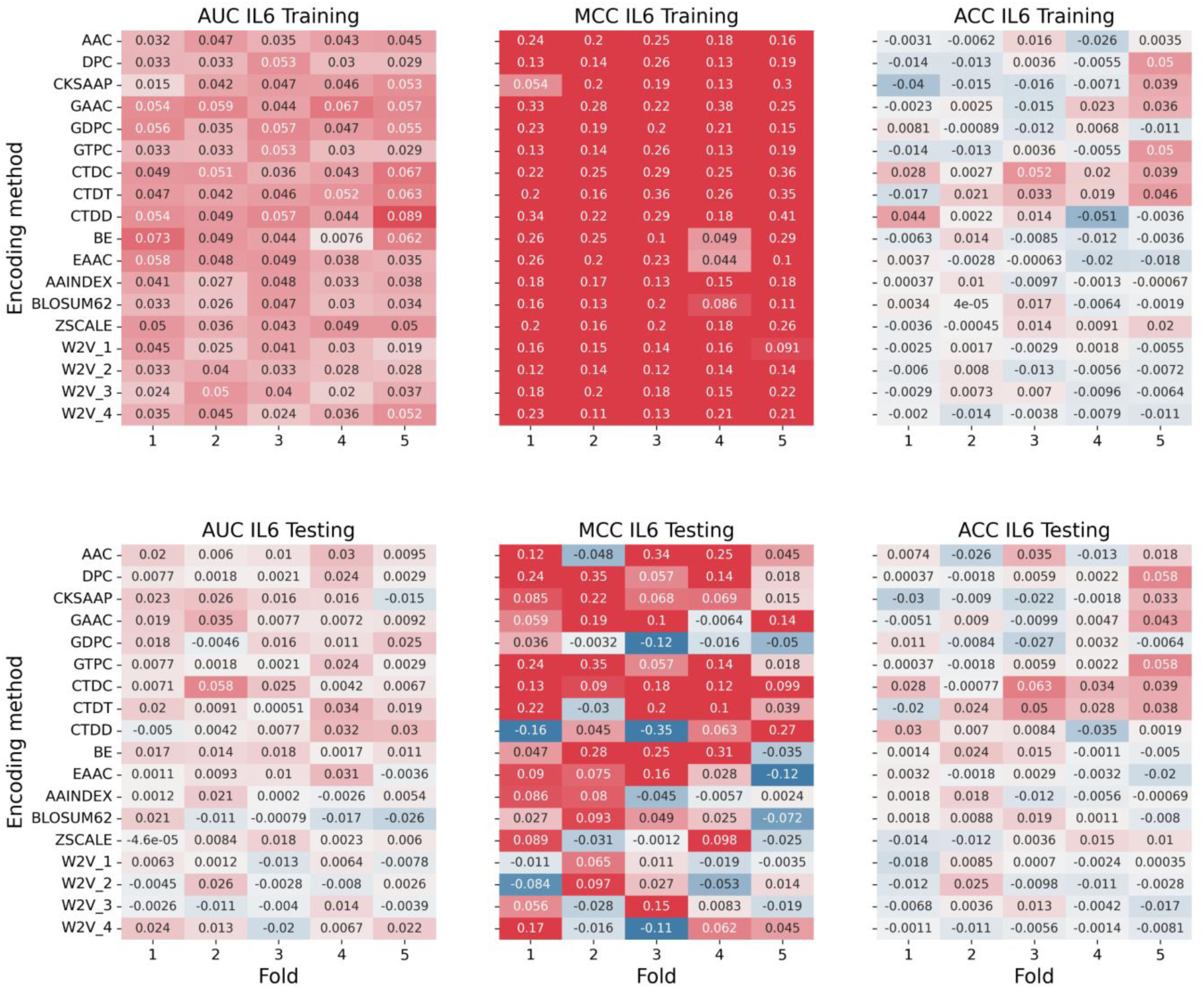
PIRs of AUC, MCC, and ACC for 18 baseline classifiers trained by the augmented training dataset with GAN-generated IL-6-inducing peptides via the stratified 5-fold CV with cluster partitioning. The optimal parameter settings of GDA-GAN were a cutoff of 0.7, an AR of 0.25, and a PT of 0.5.

**Figure 8.**
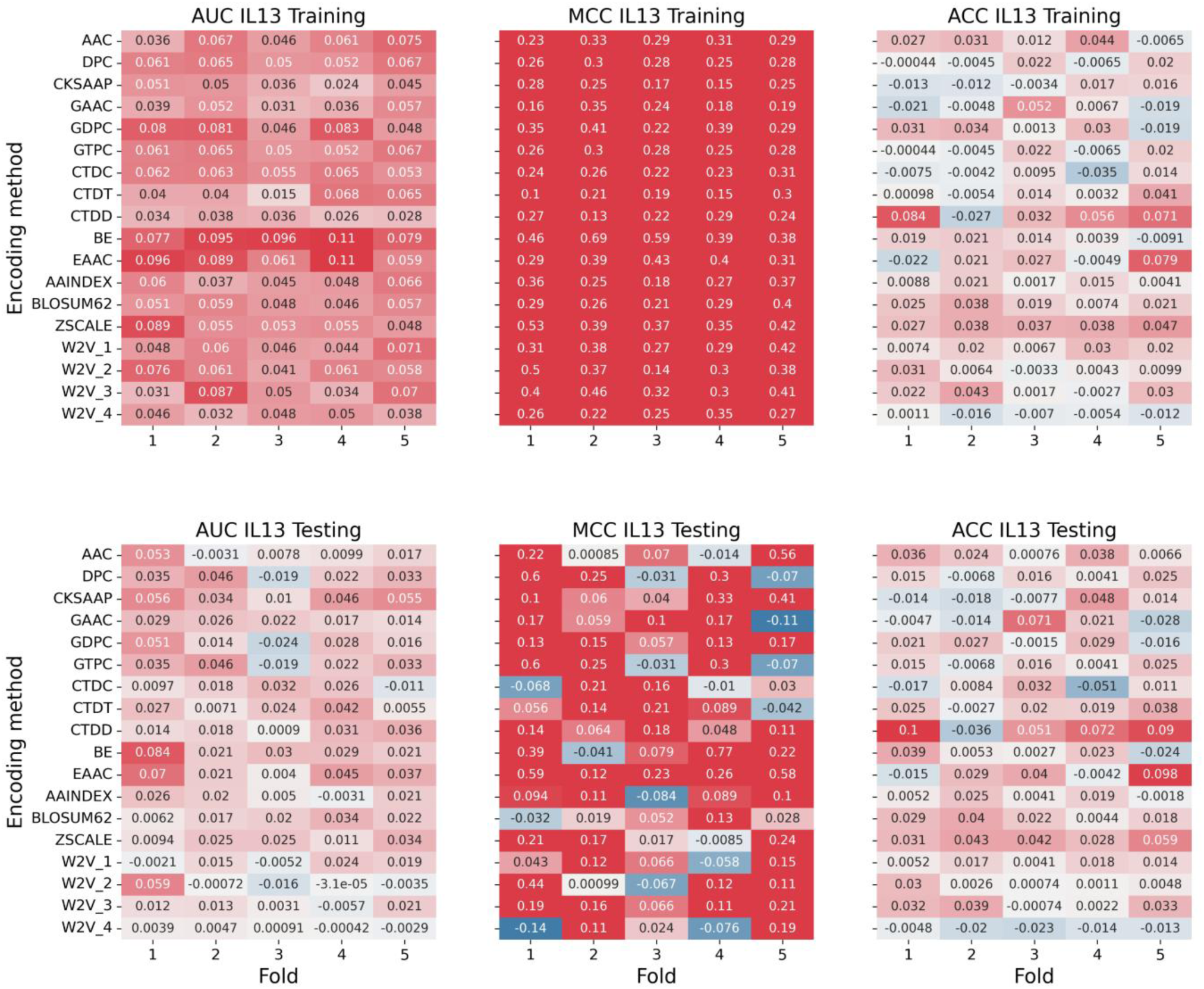
PIRs of AUC, MCC, and ACC for 18 baseline classifiers trained by the augmented training dataset with GAN-generated IL-13-inducing peptides via the stratified 5-fold CV with cluster partitioning. The optimal parameter settings of GDA-GAN were a cutoff of 0.7, an AR of 0.25, and a PT of 0.5.

To evaluate the generalizability of the GDA method, it was incorporated into SOTA models (PredIL6, PredIL13), termed GDA-Preds, for the identification of IL-6- and IL-13-inducing peptides on their benchmark datasets, as shown in **Table 4**. For PredIL6, the incorporation of GDA resulted in improvements of 1.7% in AUC, 2.1% in MCC, and 0.2% in ACC. Similarly, for PredIL13, GDA achieved increases of 0.8% in AUC, 24.3% in MCC, and 0.4% in ACC. These results demonstrate that GDA-Preds enhance the prediction performance of the PredIL6 and PredIL13 for IL6- and IL13-inducing peptides, respectively. To further confirm its effectiveness, GDA was applied to the PepNet predictor. The integration of GDA improved the performance for IL6- and IL13-inducing peptides, with respective increases of 5.3% and 1.3% in AUC, 6.8% and 19.9% in MCC, and 5.3% and 1.4% in ACC.

**Table 4.** GDA-enhanced prediction performances of SOTAs for identifying IL6- and IL13-inducing peptides on the benchmark test dataset. Two SOTA models were employed to predict AIPs with and without applying GDA. GDA is performed using GANs with a cutoff of 0.7, a PT of 0.5, and an AR of 0.25.

| Peptide | Classifier | DA | SEN | SPE | PRE | ACC | MCC | AUC |
| --- | --- | --- | --- | --- | --- | --- | --- | --- |
| IL6-inducing peptide | PredIL6 | **GDA** | **0.458** | 0.987 | 0.811 | **0.929** | **0.575** | **0.882** |
|  |  | SMOTE | 0.332 | 0.991 | 0.855 | 0.919 | 0.490 | 0.868 |
|  |  | KDE | 0.405 | **0.993** | **0.872** | 0.929 | 0.565 | 0.868 |
|  |  | w/o | 0.438 | 0.987 | 0.813 | 0.928 | 0.563 | 0.867 |
|  | PepNet | **GDA** | 0.411 | **0.984** | **0.759** | **0.921** | **0.522** | **0.873** |
|  |  | SMOTE | 0.359 | 0.984 | 0.751 | 0.916 | 0.474 | 0.857 |
|  |  | KDE | 0.422 | 0.981 | 0.748 | 0.920 | 0.522 | 0.871 |
|  |  | w/o | **0.468** | 0.962 | 0.632 | 0.908 | 0.488 | 0.829 |
| IL13-inducing peptide | PredIL13 | **GDA** | **0.273** | 0.987 | 0.729 | **0.918** | **0.411** | **0.889** |
|  |  | SMOTE | 0.257 | 0.990 | 0.760 | 0.918 | 0.405 | 0.888 |
|  |  | KDE | 0.289 | 0.986 | 0.732 | 0.918 | 0.419 | 0.872 |
|  |  | w/o | 0.146 | **0.997** | **0.852** | 0.914 | 0.330 | 0.880 |
|  | PepNet | **GDA** | **0.352** | **0.982** | **0.676** | **0.920** | **0.451** | **0.816** |
|  |  | SMOTE | 0.305 | 0.970 | 0.539 | 0.905 | 0.357 | 0.804 |
|  |  | KDE | 0.317 | 0.976 | 0.590 | 0.911 | 0.389 | 0.800 |
|  |  | w/o | 0.308 | 0.973 | 0.586 | 0.908 | 0.376 | 0.804 |

For IL-6-inducing peptides, applying the conventional data augmentation methods (SMOTE and KDE) to PredIL6 led to improvements in SPE, PRE, and MCC relative to the control method, although these improvements were smaller than those obtained with GDA. Similarly, applying SMOTE and KDE to PepNet enhanced SPE, PRE, ACC, and AUC, but again to a lesser extent than GDA. For IL13-inducing peptides, application of SMOTE and KDE to PredIL13 increased ACC and MCC compared to the control method, yet the gains remained below those achieved with GDA. Applying SMOTE and KDE to PepNet yielded minimal or no improvement. Overall, these results demonstrate that the GDA-Preds consistently enhances the predictive performance of the SOTA models across the two peptide datasets, surpassing the effectiveness of conventional data augmentation approaches.

## 4. Discussion and conclusion

Based on a case study-oriented proof-of-concept framework, we proposed the GDA method and assessed its feasibility and generalizability using a moderately sized dataset of AIPs as an interpretable example. The AIP dataset provides sufficient data to optimize the key parameters governing GDA performance, whereas such optimization is often challenging with smaller training sets. There are four key parameters: a type of generative model employed (GAN, DM, or VAE), a sequence identity cutoff of NW-RR, a PT applied to selecting generated peptides, and an AR defining the proportion of generated peptides added to the training dataset. The AIPs-optimized GDA was subsequently applied to two case studies: the identification of IL-6- and IL-13-inducing peptides, each characterized by limited dataset sizes. The application of GDA to the SOTA models (GDA-Pred) increased the prediction performance of the two cytokine-inducing peptides. Namely, GDA-Preds outperformed SOTAs for identifying two cytokine-inducing peptides. In addition, GDA surpassed the conventional data augmentation approaches (SMOTE and KDE)

Since classifiers trained with validation datasets that share sequence patterns with the training data do not accurately reflect their generalizability, we conducted a rigorous assessment of a classifier’s generalizability by employing validation datasets with distinct sequence patterns. To this end, we adopted a 5-fold CV with cluster-based partitioning. Under this validation scheme, the GDA method was developed to improve the prediction performance for two cytokine-inducing peptides.

Real functional peptides often have limited diversity or strong pattern bias and are only a tiny fraction of possible peptide space. Small peptide datasets may lead classifiers to overfit on a few sequences. Therefore, the GDA is required to produce new variants whose sequence patterns are not present in the original dataset, helping the classifier learns broader and weaker motifs than the original ones, while preserving physicochemical properties. Generated peptides are required to fill in gaps between real and fake sequences, smoothing the decision boundary and improving generalization to unseen peptides. In GANs, competition of a generator and a discriminator makes it possible to explore and design peptide sequence space. GANs are generally prone to mode collapse, causing loss of diversity or focusing on dominant pattern and vanishing gradients. DMs learn the data distribution through a multi-step denoising process, gradually refining noise into a peptide sequence. This iterative refinement allows DMs to preserve detailed motifs and structural or physicochemical features. VAEs learn a structured, continuous latent space where similar peptides are located near each other, allowing interpolation between peptides and smooth exploration of sequence space. VAEs are likely to generate average-like or blurred samples. In this study, regardless of commonly noted advantages and disadvantages, GANs outperformed DMs and VAEs, suggesting that GANs generate realistic, diverse distributions of peptide sequences. Their adversarial training framework could force the generator to closely mimic the real data distribution, while capturing diversity.

The prediction performance of GDA-based baseline models remained robust when a cutoff of NW-RR was varied between 0.6 and 0.9. Therefore, a cutoff of 0.7 was adopted for subsequent analyses. Removing sequence redundancy using NW is a crucial preprocessing step when training generative AI models for peptide design. Without any redundancy reduction, the model may simply memorize highly similar sequences rather than learning meaningful sequence–function relationships, resulting in overfitting. By filtering out duplicate and near-duplicate peptides, NW-RR enforces sequence diversity, compelling the model to capture generalizable patterns and enhancing its capacity to generate truly novel sequences. A PT of 0.5 was identified as optimal. The PT serves as a critical parameter that balances sequence diversity and convergence: lower PT values favor the generation of more diverse peptide sequences, whereas higher PT values restrict sequence variability, yielding more homogeneous outputs. Similarly, an AR of 0.25 was determined to be optimal. The AR governs the extent of GDA applied to the training dataset. Although higher AR values improved model performance on the training set, they adversely affected test performance, suggesting that excessive augmentation can induce overfitting and diminish generalizability.

The size of the training dataset is a critical factor influencing the effectiveness of GDA. When the peptide dataset is too small, GDA is prone to overfitting due to the limited sequence diversity. Conversely, when the original dataset is sufficiently large and diverse, it already encompasses most of the relevant sequence patterns. In such cases, GDA provides minimal additional benefit, as the sequence space is effectively saturated.

The three generative AI models have a lot of tuning parameters (**Table 2**). GANs involve tuning parameters such as the deep network architecture, latent vector dimension, learning rate, and discriminator update frequency. DMs involve noise schedule and number of diffusion steps, and sampling parameters. VAEs involve latent dimension, KL divergence weights. Since optimization of these parameters is an extremely time-consuming task due to calculation complexity, we did not intensively optimize the models in this study. Extensive optimization of the generative models will be explored in future work.

## Author contribution

Conceptualization: Hiroyuki Kurata. Data curation: Md. Harun-Or-Roshid. Formal analysis: Hiroyuki Kurata, Md. Harun-Or-Roshid, Kazuhiro Maeda. Funding acquisition: Hiroyuki Kurata. Investigation: Hiroyuki Kurata, Md. Harun-Or-Roshid. Methodology: Hiroyuki Kurata, Hiroto Tsuruta, Soyogu Shigetomi, Md. Harun-Or-Roshid. Project administration: Hiroyuki Kurata. Resources: Md. Harun-Or-Roshid. Software: Hiroyuki Kurata, Hiroto Tsuruta, Soyogu Shigetomi, Md. Harun-Or-Roshid. Supervision: Hiroyuki Kurata. Validation: Md. Harun-Or-Roshid Kazuhiro Maeda. Writing-original draft: Hiroyuki Kurata. Writing- review & editing: Md. Harun-Or-Roshid, Kazuhiro Maeda.

## Declarations

While preparing this work, the authors used ChatGPT based on the GPT-5.1 architecture to improve the readability. After using this tool, the authors reviewed and edited the content as needed and take full responsibility for the content of the published article.

## Ethics statement

The authors declare that the current study did not involve any experiments on humans or animals. All computational analyses were conducted using publicly available data and software tools, and no ethical approval was required. The research complies with the ethical standards of scientific integrity, transparency, and reproducibility. All authors have read and approved the manuscript and agree with its submission.

## Data availability

The programs of GDA and GDA-Pred are freely available at https://github.com/kuratahiroyuki/GDAPred.

