## Supplemental Information for "GDA-Pred: Generative AI-Driven Data Augmentation for Improved Prediction of IL-6 and IL-13 Inducing Peptides"

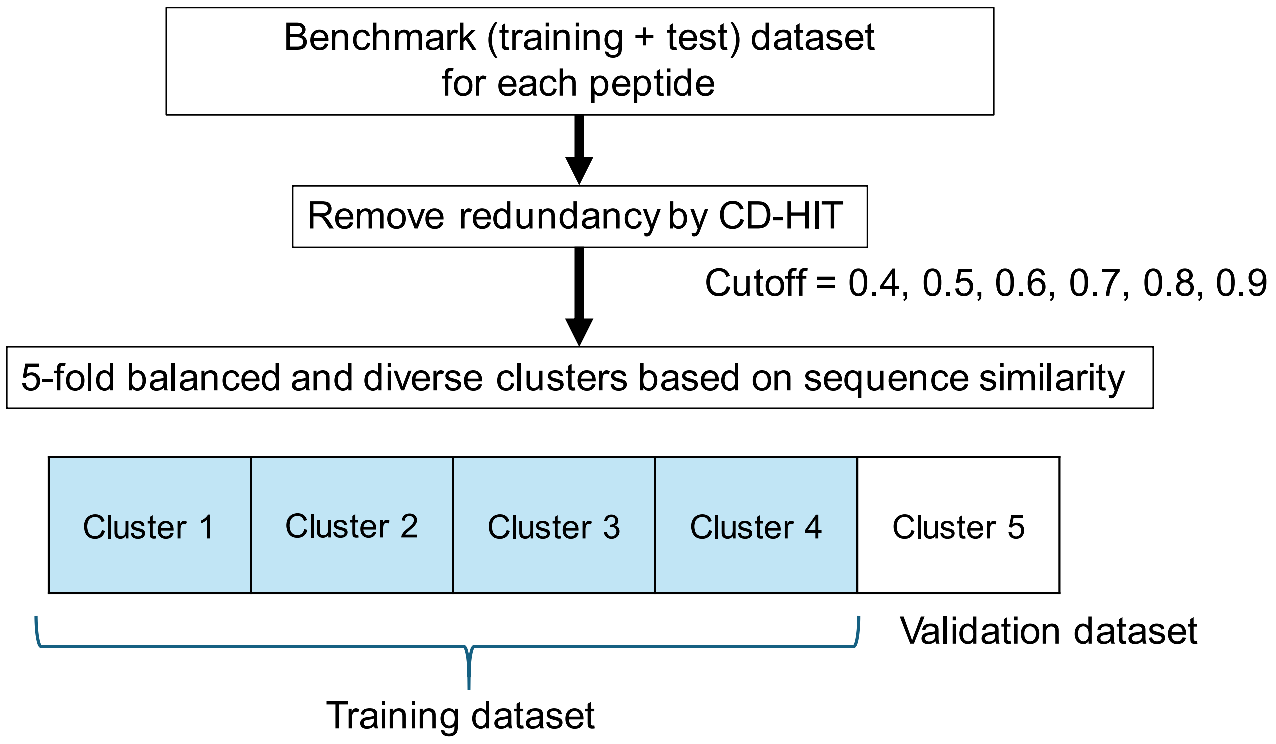


Figure S1 Stratified 5-fold CV with cluster-based partitioning for rigorously testing the generalizability of predictors.

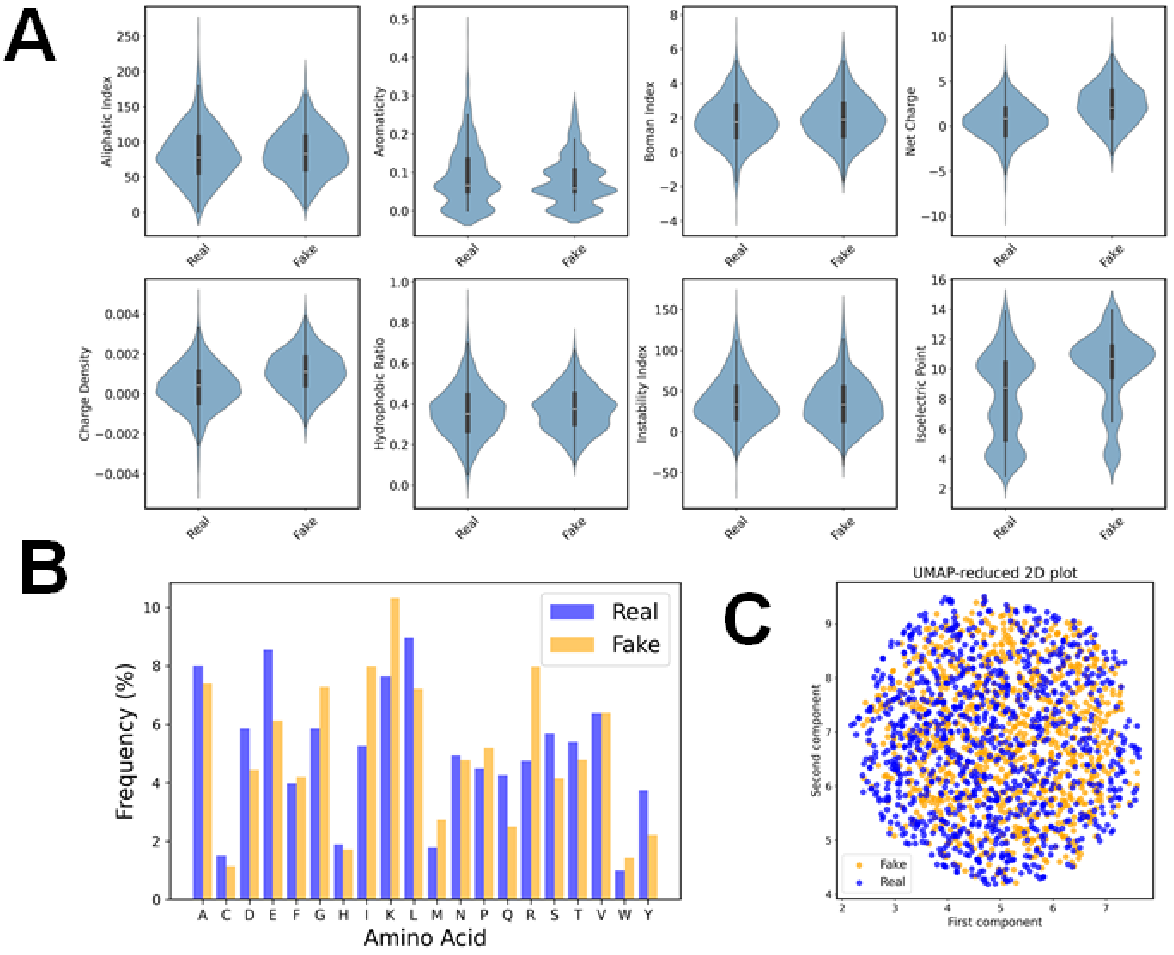


Figure S2 Evaluation of the quality of AIP fake peptide sequences that DMs generate from the real, positive peptides of the training dataset. The fake peptides correspond to the generated peptides. The fake AIPs are generated from the non-redundant training dataset built with NW-RR with a sequence identity cutoff of 0.7. (A) Physicochemical properties. (B) Frequency distribution of amino acid residues. (C) UMAP distributions of the encoding vectors of real and fake peptide sequences.

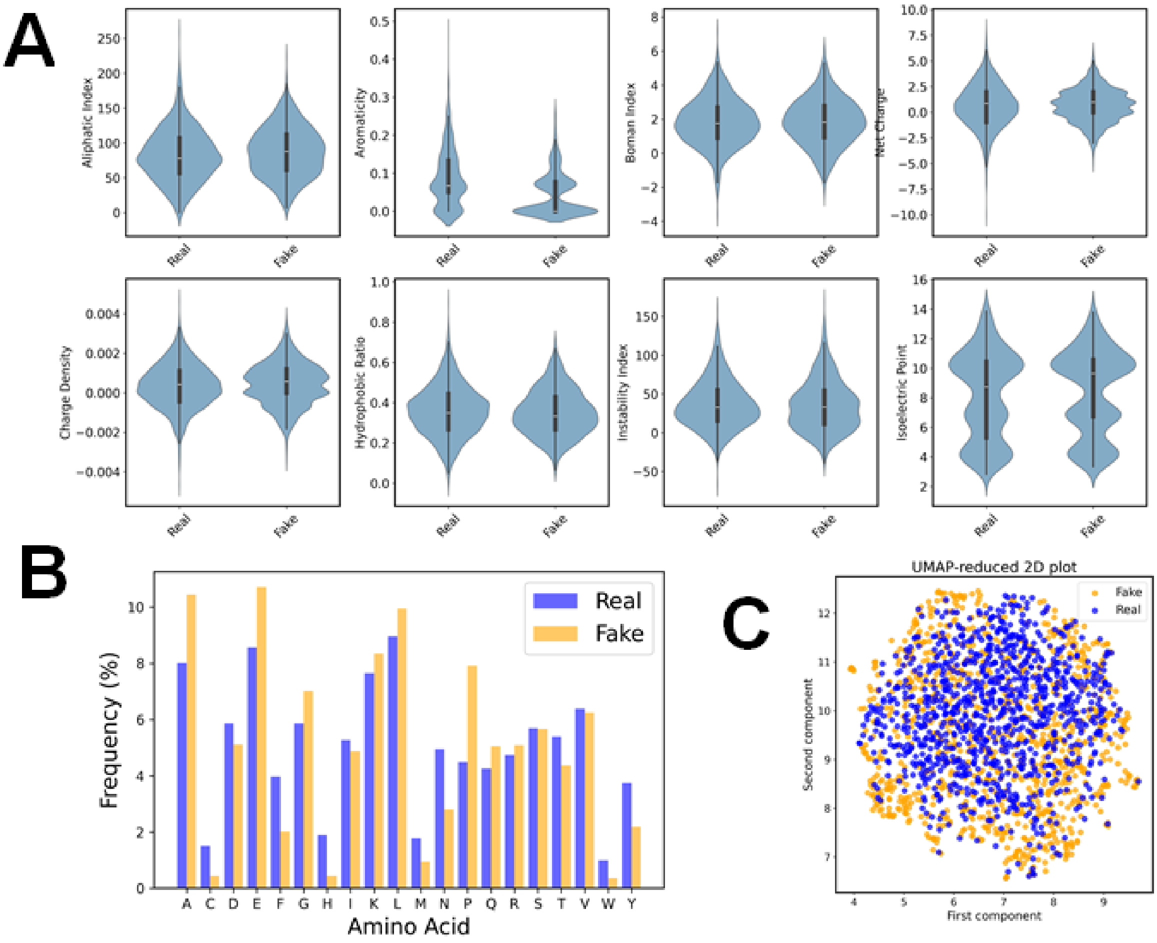


Figure S3 Evaluation of the quality of AIP fake peptide sequences that VAEs generate from the real, positive peptides of the training dataset. The fake peptides correspond to the generated peptides. The fake AIPs are generated from the non-redundant training dataset built with NW-RR with a sequence identity cutoff of 0.7. (A) Physicochemical properties. (B) Frequency distribution of amino acid residues. (C) UMAP distributions of the encoding vectors of real and fake peptide sequences.

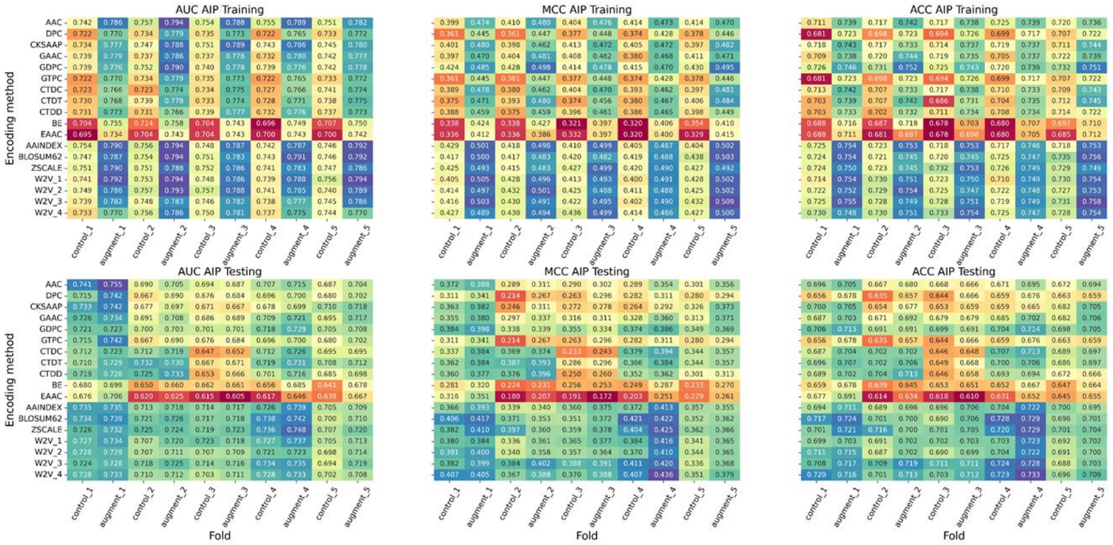


Figure S4 Prediction performance of 18 baseline classifiers trained by the augmented training dataset with GAN-generated AIPs via the stratified 5-fold CV with cluster partitioning. Fake AIP sequences were generated using GANs from a non-redundant training dataset built by CD-HIT with a sequence identity cutoff of 0.7. The generated peptides were filtered using an LGBM classifier with BLOSUM62 encoding at a PT of 0.5, and the selected sequences were added to the training dataset with an AR of 0.25.

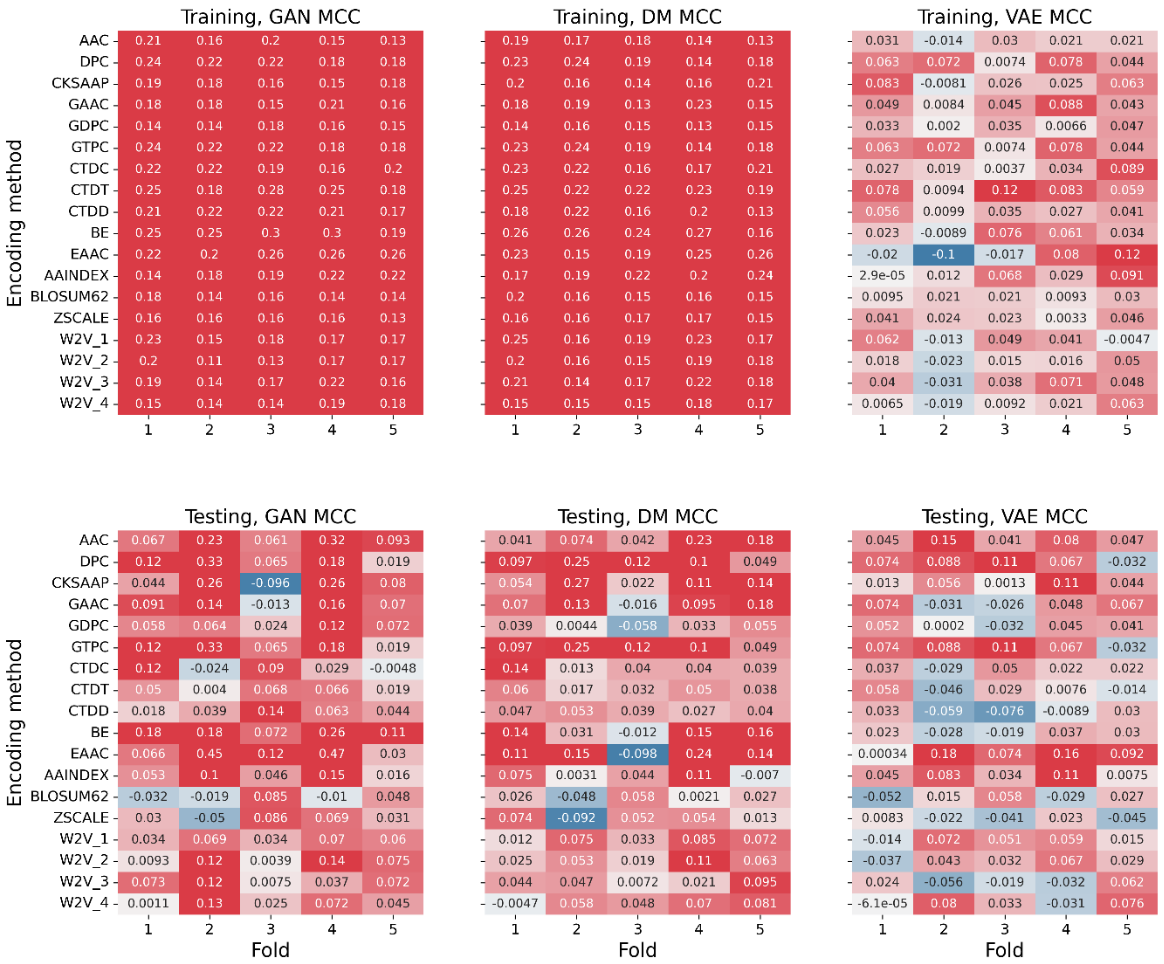


Figure S5 Comparison among the three generative AIs (GAN, DM, VAE) via stratified 5-fold CV with cluster-based partitioning, based on PIRs of MCC for 18 baseline classifiers trained by the augmented training dataset with three generative AIs-generated AIPs. Fake AIP sequences were generated using GANs from a non-redundant training dataset constructed by NW with a cutoff of 0.7. The generated peptides were selected at a PT of 0.5 using an LGBM classifier with the BLOSUM62 encoding and subsequently added to the training dataset with an AR of 0.25.

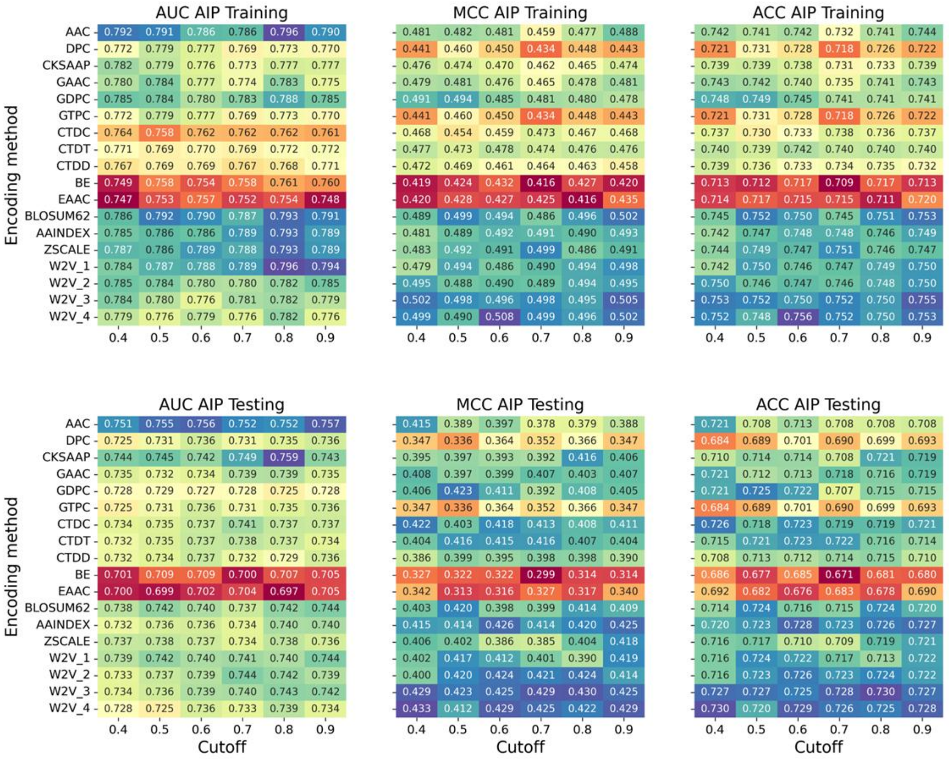


Figure S6 Effect of a sequence identity cutoff on AUC, MCC, and ACC for 18 baseline classifiers trained with the augmented training dataset with GAN-generated AIPs via the stratified 5-fold CV with cluster partitioning. Fake AIPs are generated from non-redundant training datasets built via NW with a cutoff of 0.5, 0.6, 0.7, 0.8 and 0.9. The generated peptides are selected at a PT of 0.5 by an LGBM classifier with BLOSUM62 encoding, and then added to the training dataset with an AR of 0.25.

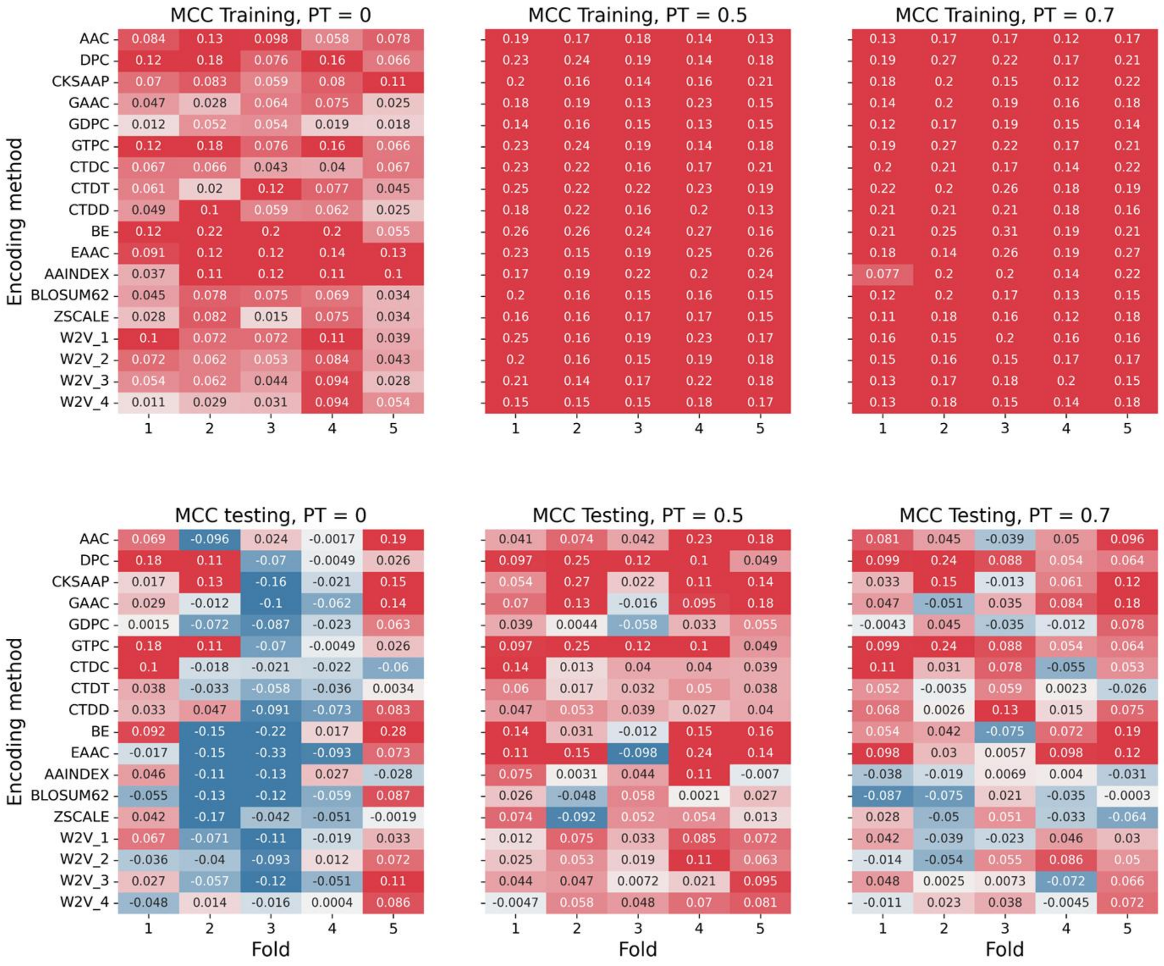


Figure S7 Effect of a PT on the PIRs of AUC, MCC, and ACC for 18 baseline classifiers trained by the augmented training dataset with GAN-generated AIPs via the stratified 5-fold CV with cluster partitioning. Fake AIP sequences were generated using GANs from a non-redundant training dataset constructed by NW with a cutoff of 0.7. The generated peptides were selected at PTs of 0, 0.5, and 0.7 using an LGBM classifier with the BLOSUM62 encoding and subsequently added to the training dataset with an AR of 0.25.

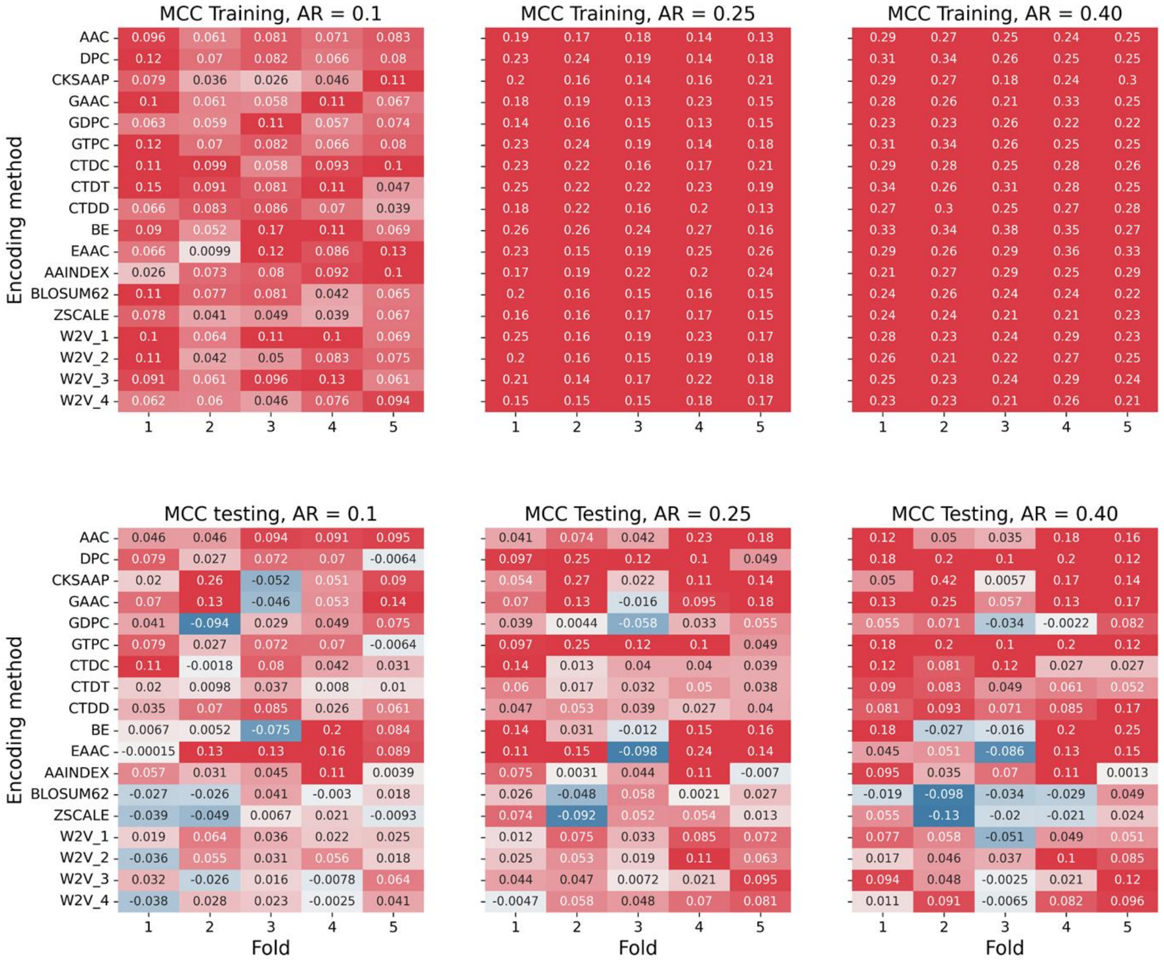


Figure S8 Effect of an AR on the PIRs of AUC, MCC, and ACC for 18 baseline classifiers trained by the augmented training dataset with GAN-generated AIPs via the stratified 5-fold CV with cluster partitioning. Fake AIP sequences were generated using GANs from a non-redundant training dataset constructed by NW with a cutoff of 0.7. The generated peptides were selected at a PT of 0.5 using an LGBM classifier with the BLOSUM62 encoding and subsequently added to the training dataset with ARs of 0.1, 0.25, and 0.4.
